# Longitudinal effects of using and discontinuing CNS medications on cognitive functioning

**DOI:** 10.1101/2021.09.13.460082

**Authors:** Elise Koch, Kristina Johnell, Karolina Kauppi

**Author notes:** Address correspondence to Dr. Elise Koch, NORMENT Centre, Oslo University and Oslo University Hospital, 0424 Oslo, Norway.

## Abstract

**Purpose:** To investigate the longitudinal effect of using and discontinuing central nervous system (CNS) medications on cognitive performance.

**Methods:** Using longitudinal cognitive data from healthy adults aged 25-100 years (N = 2,188) from four test waves five years apart, we investigated both the link between use of CNS medications (opioids, antidepressants, and anxiolytics, hypnotics and sedatives) on cognitive task performance (episodic memory, semantic memory, visuospatial ability) across 15 years, and the effect of discontinuing these medications in linear mixed effects models.

**Results:** We found that opioid use was associated with decline in visuospatial ability, whereas antidepressant use was associated with decline in semantic memory over 15 years. A link between drug discontinuation and cognitive improvement was seen for opioids, antidepressants as well as for anxiolytics, hypnotics and sedatives.

**Conclusions:** Although our results may be confounded by subjacent conditions, they suggest that long-term use of CNS medications may have domain-specific negative effects on cognitive performance over time, whereas the discontinuation of these medications may partly reverse these effects. These results open up for future studies that address subjacent conditions on cognition to develop a more complete understanding of the cognitive effects of CNS medications.

**Key points:**

- Opioid use was associated with decline in visuospatial ability, and individuals discontinuing using opioids showed improvement in visuospatial ability compared to individuals continuing using opioids.
- Antidepressant use was associated with decline in semantic memory, and individuals discontinuing using antidepressants showed improvement in semantic memory compared to individuals continuing using antidepressants.
- For anxiolytics, hypnotics and sedatives there was no difference between continued users and non-users, but drug discontinuation was associated with more positive cognitive development both in episodic memory and visuospatial ability.

## Introduction

Medications acting on the central nervous system (CNS) frequently cause adverse effects, including problems with mobility, falls, and cognition in older individuals.^1–3^ Results from longitudinal studies investigating cognitive effects of CNS medication use suggest that CNS medications may accelerate cognitive decline in older adults,^4–7^ and it has been suggested that cognitive problems related to CNS medication use may be reversed by adjusting or discontinuing these medications.^8^ Accelerated cognitive decline as well as increased risk of developing dementia has been mostly related to CNS medications with strong anticholinergic properties^8,9^ including opioids, antidepressants, and antipsychotics, but also benzodiazepines or related drugs (anxiolytics, hypnotics, and sedatives) as well as antiepileptics.^4–7^

A recent systematic review^10^ on the impact of opioid use on cognition in older adults showed that both improvements and impairments of cognition have been observed, especially in studies with higher mean opioid doses. Whereas most of these studies demonstrated no effect of opioid use on task performance in any cognitive domain, other studies showed that attention, language, orientation, psychomotor function, verbal working memory, and episodic memory were worsened, especially at high mean doses of opioids.^10^ It has also been shown that individuals using opioids at high doses had slightly higher dementia risk than individuals with little or no use of opioids.^11^ However, these results may reflect cognitive effects of chronic pain,^11^ and it has been suggested that in patients taking opioids for chronic pain, severity and psychological measures are more predictive of cognitive decline that opioid use.^12^ For antidepressants, findings from studies investigating cognitive effects are mixed, with cognitive impairments, no effects, and pro-cognitive effects having been reported in relation to antidepressant use.^13–15^ Cognitive impairments have mostly been related to the use of tricyclic antidepressants (TCAs), which is probably related to their anticholinergic effects, whereas serotonin selective reuptake inhibitors (SSRIs) have little if any anticholinergic properties and less cognitive impairment in depressed patients.^14^ Evidence suggests that SSRIs may in fact improve cognitive functions in depressed patients^13,14,16^ through mechanisms separate from their antidepressant effects.^14^ However, depressive symptoms have known negative effects on cognition,^17^ and in non-depressed individuals, antidepressants did not significantly affect cognitive functioning in any direction.^16^ Moreover, a longitudinal study showed that cognition declined in individuals taking antidepressants at the same rate as those not taking antidepressants when adjusting for depressive symptoms.^18^ Benzodiazepines and related drugs enhance the effect of the inhibitory neurotransmitter gamma-aminobutyric acid (GABA)^19^ that may modulate cognitive performance,^20^ and their acute negative effects on cognitive functioning are well established.^21–23^ Moreover, there is mounting evidence of an association between long-term use of benzodiazepines and impairments in a range of neuropsychological functions, and some evidence suggests a higher risk of decline in cognitive functioning of various cognitive domains as a result of long-term benzodiazepine use.^24^ Benzodiazepine use has also been found to be strongly associated with an increased risk of dementia.^25^ Since insomnia and anxiety are prodromal symptoms of dementia, it is difficult to determine whether this association arises from the treatment of those early symptoms prior to diagnosis or if benzodiazepines and related drugs are causing cognitive decline and dementia.^26^

The longitudinal effects of CNS medications on cognitive performance are still not fully understood, and discrepancies between studies are likely due to methodological differences and inconsistencies regarding the cognitive domains affected.^27^ In the present study, we investigated if use of CNS medications (i.e. opioids, antidepressants, anxiolytics, hypnotics and sedatives) is associated with cognitive functioning (episodic memory, semantic memory, visuospatial ability) over time. Using longitudinal cognitive data from healthy adults aged 25-100 years, we investigated both the association between CNS medication use, and the association between discontinuing these medications and cognitive task performance.

## Methods

### Participants

Data in the present study come from the longitudinal population-based Betula Prospective Cohort Study on memory, health and aging, conducted in Umeå, Sweden,^28,29^ including measurements of cognitive functions from four test waves (T3-T6) five years apart, with a total follow-up period of 15 years. Exclusion criteria were dementia and known severe neurologic or psychiatric disease. From five cohorts (S1 and S3 (T3-T6), S2 and S4 (T3), and S6 (T5-T6)), data on drug use was available for 2,895 individuals. We excluded those who developed dementia (N = 394) and those who were included at T5 (S6) and only had one visit (N = 292) as well as those with missing cognitive data (N = 21) resulting in 2,188 individuals aged 25-100 years (with an average of 1.986 visits, ranging from 1-4) included in the current study. Dementia diagnosis was done by a geropsychiatrist based on the DSM-IV criteria^30^ (the diagnosis procedure has been described elsewhere).^28,31,32^ The research was approved by the regional ethical review board at Umeå University (EPN), and all participants gave written informed consent.

### Use of CNS medications

Data on self-reported drug use was retrieved at each test wave, and included drugs used both regularly and as needed. CNS medications were classified according to the Anatomical Therapeutic Chemical (ATC) classification system. The groups of medications defined as those having CNS effects and used in our analyses were as follows: Opioids (ATC code N02A), antidepressants (ATC code N06A), anxiolytics (ATC code N05B), hypnotics and sedatives (ATC code N05C). For each CNS medication user and time-point, 1-3 matched controls were chosen. Controls were matched on sex, age, years of education, test wave as well as sample, and chosen among individuals not taking any CNS medications, which on top of the above-mentioned also included anticonvulsants (ATC code N03A), anti-Parkinson drugs (ATC codes N04A and N04B), and antipsychotics (ATC code N05A). Matching was done separately for individuals that only had one visit (at T3) and individuals that had more than one visit to also choose controls that either had one or more visits. For our analyses, CNS medication use (used regularly or as needed versus no use) was coded as three dichotomized variables: Opioids (ATC code N02A), antidepressants (ATC code N06A), and anxiolytics, hypnotics and sedatives (ATC codes N05B and N05C). To investigate longitudinal effects of using CNS medications, our analyses focused on participants who used CNS medications at both baseline and all their follow-up examinations. To investigate the effects of discontinuing CNS medications, two consecutive time-points were analyzed, where individuals were taking the drug at the first time-point but not at the following time-point. Individuals discontinuing CNS medications were analyzed both in relation to matched controls not using any CNS medication at both time-points (matched on sex, age, years of education, test wave as well as sample) as well as in relation to individuals continuing using CNS medications (matched on years of education, test wave, and sample) to investigate the effects of discontinuing CNS medications in relation to both groups.

### Cognitive tests

The analyses were based on z-transformed tests of visuospatial ability, episodic memory and semantic memory. For episodic memory, a composite score was calculated as the sum of z-transformed tests of episodic memory (two tests of free oral recall of verb-noun sentences, two tests of category-cued recall of nouns from the sentence recall, and one test of free recall of presented nouns). For semantic memory, a composite score was calculated as the sum of z-transformed tests of semantic knowledge/verbal fluency (three tests of verbal generation of as many words as possible during 1 min: words that begin with A; Words that begin with M, exactly 5 letters; professions that begin with B). Visuospatial ability was measured with the block design task from the Wechsler Adult Intelligence Scale (WAIS-R).^33^ For details about cognitive tasks see Nilsson et al. 2004.^28^

### Statistical analyses

To examine if CNS drug use is associated with level and change in cognitive measures over 15 years, we performed linear mixed-effect models including individuals taking CNS mediations at all their visits and matched controls not taking any CNS medication at any of their visits. These models were fitted in R using the lme4 function available through the lme4 package. P-values were estimated based on the Satterthwaite approximations implemented in the lme4Test package. The models included the dichotomized drug variable as covariate of interest and the following covariates of no interest: age, age^2^, sex, and number of visits between T3 and T6 (to control for potential drop-out effects). Time from inclusion (years) was used as time-scale to represent slope, and interaction with time was allowed for all covariates. The models also included random subject-specific intercepts. The same models were used to examine if discontinuing using CNS medications is associated with level or change in cognitive measures, but these models included only two time-points, where individuals using CNS medications were taking the drug at the first time-point but not at the following time-point and either matched controls not using any CNS medication at both time-points or matched controls using the same CNS medication at both time-points. Levene’s test was used to assess the equality of variance in baseline cognitive test performance in users and non-users of CNS medications. All statistical analyses were performed in R version 4.0.3.

## Results

Descriptive statistics for each test occasion (T3-T6) are shown in **Table 1**. Descriptive statistics per group (individuals using CNS medications, individuals discontinuing CNS medications, and individuals not using any CNS mediations) are shown in **Table 2**. Based on a cognition composite score that was calculated as the sum of z-transformed tests of visuospatial ability, episodic memory and semantic memory, the equality of variance in baseline cognitive test performance did not differ between users and non-users of opioids (f = 0.139, p-value = 0.704), antidepressants (f = 0.597, p-value = 0.440), and anxiolytics, hypnotics and sedatives (f = 0.611, p-value = 0.435). **Figure 1** shows the cognitive test performance of individual cognitive tests across 15 years separately for individuals not using any CNS medications and individuals using opioids (**Figure 1A**), antidepressants (**Figure 1B**), and anxiolytics, hypnotics and sedatives (**Figure 1C**).

**Table 1:** Demographic, cognition-related and drug-related variables of participants at each test occasion

|  | **T3**<br>(N =<br>**2,126**) | **T4**<br>(N =<br>**919**) | **T5**<br>N =<br>**753**) | **T6**<br>(N =<br>**496**) |
| --- | --- | --- | --- | --- |
| **Age**, Mean (SD) | 60.97<br>(13.39) | 64.55<br>(11.06) | 65.44<br>(11.47) | 67.81<br>(10.79) |
| **Sex**, N females (males) | 1,092<br>(1,034) | 490<br>(429) | 403<br>(350) | 261<br>(235) |
| **Years of education**, Mean (SD) | 10.93<br>(4.20) | 11.65<br>(4.27) | 12.41<br>(4.27) | 12.93<br>(4.37) |
| **Cognitive task performance**, Mean (SD) |  |  |  |  |
| Episodic memory | 35.91<br>(10.84) | 38.02<br>(10.51) | 38.44<br>(10.58) | 38.86<br>(10.52) |
| Semantic memory | 21.40<br>(8.15) | 23.89<br>(8.61) | 23.68<br>(8.27) | 24.50<br>(8.66) |
| Visuospatial ability | 26.56<br>(10.88) | 27.35<br>(9.88) | 27.74<br>(10.07) | 27.18<br>(9.81) |
| **CNS medications**, N users (non-users) |  |  |  |  |
| Opioids | 70<br>(2,056) | 15<br>(904) | 10<br>(743) | 5<br>(491) |
| Antidepressants | 23<br>(2,103) | 13<br>(906) | 10<br>(743) | 6<br>(490) |
| Anxiolytics, hypnotics and sedatives | 96<br>(2,030) | 29<br>(890) | 17<br>(736) | 6<br>(490) |

SD = standard deviation.

**Table 2:** Baseline characteristics per group

|  | **Users**<br>**continued** | **Users**<br>**discontinued** | **Non-**<br>**users** |
| --- | --- | --- | --- |
| **Opioids** | **N = 71** | **N = 34** | **N = 2083** |
| **Age**, Mean (SD) | 66.83<br>(14.64) | 61.18<br>(14.20) | 60.34<br>(13.57) |
| **Sex**, N females (males) | 41<br>(30) | 29<br>(5) | 1049<br>(1034) |
| **Years of education**, Mean (SD) | 9.49<br>(3.86) | 10.37<br>(3.60) | 11.07<br>(4.22) |
| **Task performance**, Mean (SD) |  |  |  |
| Episodic memory | 31.68<br>(10.60) | 36.59<br>(10.11) | 36.24<br>(10.89) |
| Semantic memory | 18.63<br>(7.33) | 21.26<br>(8.60) | 21.54<br>(8.14) |
| Visuospatial ability | 22.07<br>(11.00) | 27.94<br>(10.60) | 26.95<br>(10.93) |
| **Number of visits T3-T6**, Mean (SD) | 1.49 (0.89) | 3.21 (0.91) | 1.98 (1.21) |
| **Antidepressants** | **N = 24** | **N = 14** | **N = 2150** |
| **Age**, Mean (SD) | 67.92<br>(16.28) | 54.62<br>(15.00) | 60.52<br>(13.60) |
| **Sex**, N females (males) | 16<br>(8) | 7<br>(7) | 1096<br>(1054) |
| **Years of education**, Mean (SD) | 11.35<br>(3.70) | 12.32<br>(4.29) | 10.99<br>(4.21) |
| **Task performance**, Mean (SD) |  |  |  |
| Episodic memory | 34.96<br>(13.26) | 39.71<br>(11.47) | 36.09<br>(10.87) |
| Semantic memory | 22.63<br>(8.39) | 25.21<br>(7.28) | 21.40<br>(8.14) |
| Visuospatial ability | 24.88<br>(11.22) | 29.57<br>(9.55) | 26.81<br>(10.96) |
| **Number of visits T3-T6**, Mean (SD) | 2.17 (1.20) | 3.00 (0.88) | 1.98 (1.21) |
| **Anxiolytics, hypnotics, sedatives** | **N = 96** | **N = 17** | **N = 2075** |
| **Age**, Mean (SD) | 68.70<br>(13.32) | 63.82<br>(13.05) | 60.16<br>(13.56) |
| **Sex**, N females (males) | 53<br>(43) | 10<br>(7) | 1056<br>(1019) |
| **Years of education**, Mean (SD) | 10.13<br>(4.48) | 10.53<br>(4.43) | 11.05<br>(10.13) |
| **Task performance**, Mean (SD) |  |  |  |
| Episodic memory | 31.04<br>(11.07) | 37.00<br>(8.22) | 36.32<br>(10.86) |
| Semantic memory | 18.68<br>(8.60) | 20.88<br>(7.11) | 21.57<br>(8.10) |
| Visuospatial ability | 21.34<br>(11.35) | 23.82<br>(9.13) | 27.08<br>(10.88) |
| **Number of visits T3-T6**, Mean (SD) | 1.55 (0.89) | 3.06 (0.83) | 2.00 (1.22) |

SD = standard deviation.

**Figure 1:**
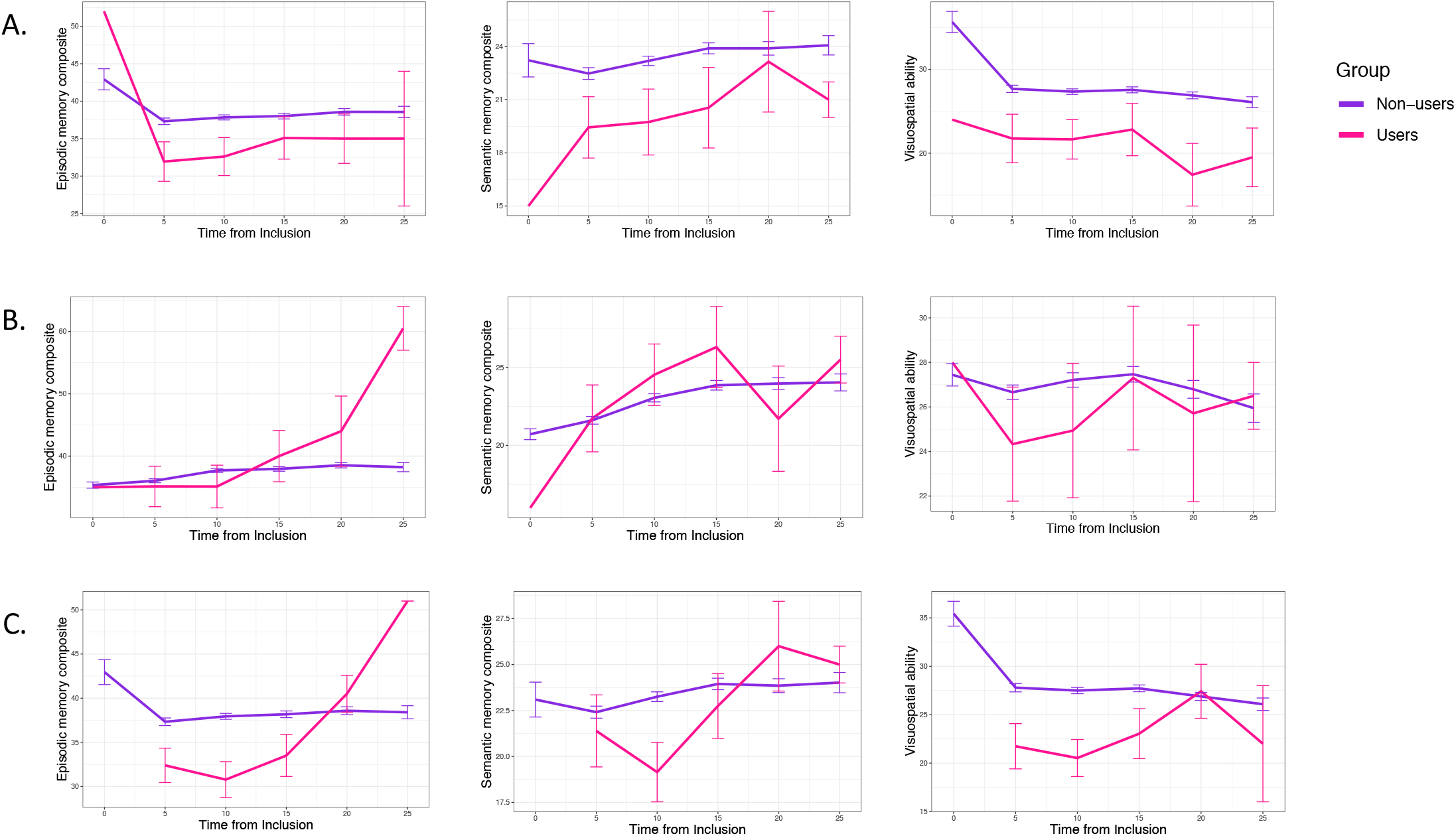
Cognitive test performance of individual cognitive tests across 15 years. Shown separately for individuals not using any CNS medications and individuals using opioids (**A**), antidepressants (**B**), anxiolytics, hypnotics and sedatives (**C**). Error bars show standard error.

### Effect of using CNS medications on cognitive performance

To analyze the effect of using CNS medications on cognitive intercept and slope, we compared individuals taking the same CNS drug at all their visits with 1-3 matched controls per individual and visit. As shown in **Table 3**, using opioids was associated with cognitive decline of visuospatial ability, whereas using antidepressants was associated with decline in semantic memory. No associations were seen for anxiolytics, hypnotics and sedatives, even at uncorrected levels (**Table 3**).

**Table 3:**
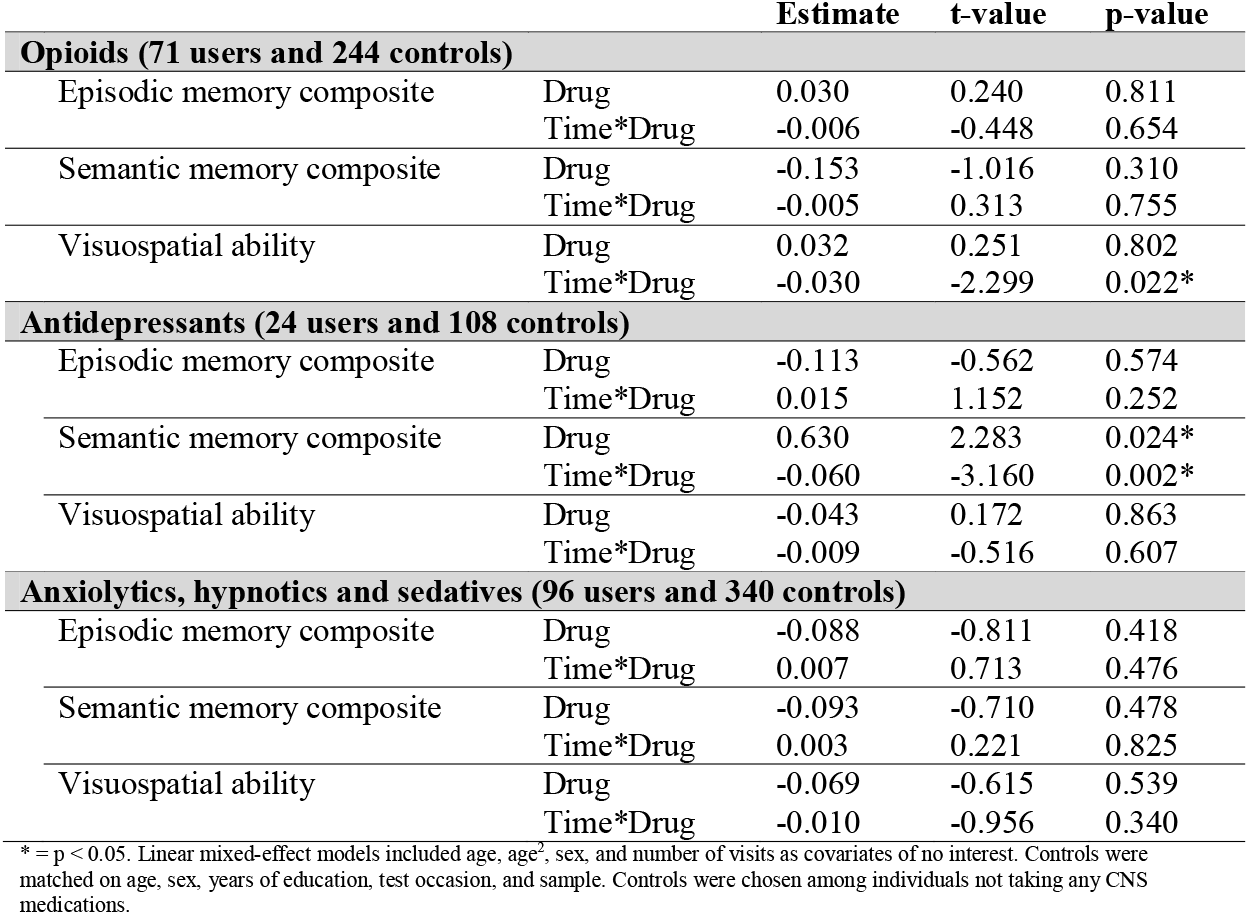
Effect of using CNS medications on intercept and slope of individual cognitive tests, including all four test occasions (T3-T6).

### Effect of discontinuing CNS medications on cognitive performance

To analyze the effect of discontinuing CNS medications on cognitive performance over time, we included the two time-points of individuals that were taking a CNS drug at one visit but not at their next visit. In relation to individuals not taking any CNS medications at both these time-points, discontinuing anxiolytics, hypnotics and sedatives was significantly associated with cognitive improvement in both episodic memory and visuospatial ability whereas discontinuing antidepressants had no significant impact on cognitive performance. Individuals discontinuing opioids significantly declined in visuospatial ability compared to individuals not using any CNS medications (**Table 4**). In relation to individuals using the same CNS medication at both time-points, discontinuation of both opioids and anxiolytics, hypnotics and sedatives was significantly associated with cognitive improvement in visuospatial ability, and there was a positive effect of discontinuing antidepressants on cognitive improvement in semantic memory (**Table 5**). The effect of discontinuing CNS medications on individual cognitive tests over time including two test occasions per individual from all four test occasions is shown in **Figure 2** for opioids (**Figure 2A**), antidepressants (**Figure 2B**), and anxiolytics, hypnotics and sedatives (**Figure 2C**) both in relation to individuals continuing using the same CNS medication and in relation to individuals not using any CNS medications at both test occasions.

**Table 4:** Effect of discontinuing CNS medications on cognitive task performance, including two test occasions from T3-T6 where individuals were taking the drug at the first time-point but not at the following time-point and matched controls not taking any CNS medications at both occasions.

|  |  | Estimate | t-value | p-value |
| --- | --- | --- | --- | --- |
| **Opioids (34 users discontinuing and 66 controls)** |  |  |  |  |
| Episodic memory composite | Drug | -0.403 | -1.349 | 0.179 |
|  | Time*Drug | 0.015 | 0.667 | 0.506 |
| Semantic memory composite | Drug | -0.164 | -0.500 | 0.618 |
|  | Time*Drug | 0.009 | 0.383 | 0.702 |
| Visuospatial ability | Drug | 0.651 | 2.444 | 0.016* |
|  | Time*Drug | -0.046 | -2.490 | 0.014* |
| **Antidepressants (14 users discontinuing and 33 controls)** |  |  |  |  |
| Episodic memory composite | Drug | -0.228 | -0.584 | 0.561 |
|  | Time*Drug | 0.010 | 0.332 | 0.741 |
| Semantic memory composite | Drug | -0.172 | -0.353 | 0.725 |
|  | Time*Drug | 0.050 | 1.291 | 0.200 |
| Visuospatial ability | Drug | -0.387 | -0.956 | 0.342 |
|  | Time*Drug | 0.020 | 0.667 | 0.507 |
| **Anxiolytics, hypnotics and sedatives (17 users discontinuing and 65 controls)** |  |  |  |  |
| Episodic memory composite | Drug | -0.447 | -1.466 | 0.144 |
|  | Time*Drug | 0.061 | 2.592 | 0.010* |
| Semantic memory composite | Drug | -0.136 | -0.399 | 0.690 |
|  | Time*Drug | 0.020 | 0.751 | 0.454 |
| Visuospatial ability | Drug | -0.611 | -2.084 | 0.039* |
|  | Time*Drug | 0.047 | 2.194 | 0.030* |

\* = $p < 0.05$ . Linear mixed-effect models included age, age<sup>2</sup>, sex, and number of visits as covariates of no interest. Controls were matched on age, sex, years of education, test occasion, and sample. Controls were chosen among individuals not taking any CNS medications.

**Table 5:** Effect of discontinuing CNS medications on cognitive task performance, including two test occasions from T3-T6 where individuals were taking the drug at the first time-point but not at the following time-point and controls taking CNS medications at both occasions.

|  |  | **Estimate** | **t-value** | **p-value** |
| --- | --- | --- | --- | --- |
| **Opioids (34 users discontinuing and 21 controls continuing)** |  |  |  |  |
| Episodic memory composite | Drug | 0.024 | 0.063 | 0.950 |
|  | Time*Drug | -0.007 | -0.294 | 0.770 |
| Semantic memory composite | Drug | 0.004 | 0.010 | 0.992 |
|  | Time*Drug | 0.006 | 0.224 | 0.823 |
| Visuospatial ability | Drug | -0.332 | -0.841 | 0.402 |
|  | Time*Drug | 0.051 | 2.066 | 0.043* |
| **Antidepressants (14 users discontinuing and 21 controls continuing)** |  |  |  |  |
| Episodic memory composite | Drug | 0.213 | 0.522 | 0.604 |
|  | Time*Drug | -0.036 | -1.313 | 0.195 |
| Semantic memory composite | Drug | -0.929 | -1.619 | 0.111 |
|  | Time*Drug | 0.085 | 2.033 | 0.047* |
| Visuospatial ability | Drug | -0.380 | -0.837 | 0.406 |
|  | Time*Drug | 0.042 | 1.304 | 0.198 |
| **Anxiolytics, hypnotics and sedatives (17 users discontinuing and 36 controls continuing)** |  |  |  |  |
| Episodic memory composite | Drug | 0.076 | 0.245 | 0.807 |
|  | Time*Drug | 0.008 | 0.320 | 0.749 |
| Semantic memory composite | Drug | -0.338 | -0.871 | 0.386 |
|  | Time*Drug | 0.012 | 0.403 | 0.688 |
| Visuospatial ability | Drug | -0.587 | -1.776 | 0.079 |
|  | Time*Drug | 0.053 | 2.316 | 0.023* |

\* = $p < 0.05$ . Linear mixed-effect models included age, age<sup>2</sup>, sex, and number of visits as covariates of no interest. Controls were matched on years of education, test occasion, and sample. Controls were chosen among individuals taking the same CNS medication at all their visits.

**Figure 2:**
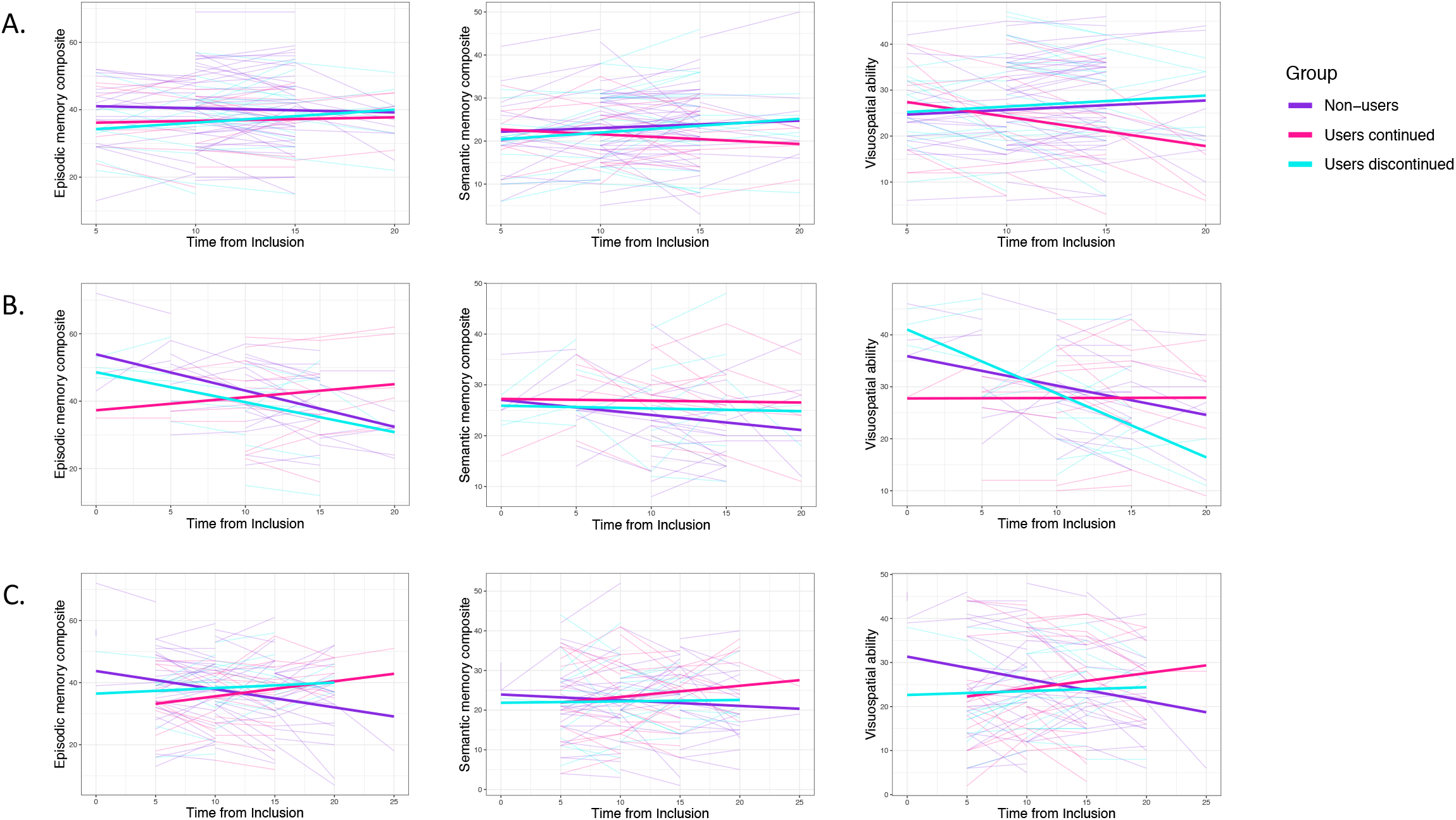
Effect of discontinuing CNS medications on cognitive performance of individual cognitive tests over time including two test occasions from T3-T6 per individual. **A**. Opioids. **B**. Antidepressants. **C**. Anxiolytics, hypnotics and sedatives. Users discontinued: Individuals that were taking a CNS drug at one visit but not at their next visit. Users continued: Matched controls taking the same CNS medication at both these test occasions. Non-users: Matched controls not taking any CNS medications at both these test occasions.

## Discussion

To our knowledge, this is the first longitudinal, population-based study investigating both the effect of using CNS medications and discontinuing these medications on cognitive task performance. Using longitudinal cognitive data from population representative adults aged 25-100 years, we found associations of CNS medication use and cognitive decline that were specific to both cognitive domains and medication class. Firstly, opioid use was associated with decline in visuospatial ability, and individuals that discontinued taking opioids had a more positive cognitive slope than individuals that continued using opioids five years later – yet more negative than those never taking opioids. Secondly, antidepressant use was associated with steeper decline in semantic memory over 15 years relative to non-users, and an effect of more positive semantic memory trajectories after drug discontinuation. Finally, for anxiolytics, hypnotics and sedatives there was no difference between continued users and non-users, but drug discontinuation was associated with more positive cognitive development both in relation to those that continued using these medications and non-users.

Consistent with previous studies, we found no differences in memory decline between individuals using opioids and matched controls.^10^ The observed greater cognitive decline in visuospatial ability in opioid users may be novel, as we are not aware of any longitudinal studies investigating the effects of opioids on cognitive decline specifically in tasks of visuospatial ability. This novel effect of cognitive decline in visuospatial ability is confirmed by our observation that individuals discontinuing using opioids show cognitive improvement in visuospatial ability compared to individuals continuing using opioids. Our observation of greater decline in visuospatial ability in individuals discontinuing using opioids compared to individuals not using opioids may indicate that the negative cognitive effects on visuospatial ability as a result of opioid use may remain to some degree after discontinuing these medications. However, we lacked measures of pain, and it has been suggested that chronic pain may be associated with impairments in cognitive functions.^34,35^ Opioids are efficient analgesics and could improve cognitive function as a result of pain reduction.^36^ On the other hand, it has been shown that opioids have negative effects on cognitive functioning also in healthy volunteers,^37^ and that opioids induce apoptosis of microglia^38,39^ and neurons.^38^ Taken together, while it remains difficult to delineate the effects of pain and opioids on cognitive functioning, our data suggest that long-term opioid use might contribute to cognitive decline in visuospatial ability.

The use of antidepressants has been associated with negative,^40^ positive,^16^ and a lack of effects on cognitive function.^18^ These results may be complicated by the potential for antidepressants to reverse the well-documented negative effect of depressive symptoms on cognition,^17,41^ together with potential direct effects of drugs. In our study, antidepressant use was associated with decline in semantic memory, but not with decline in episodic memory or visuospatial ability. However, we also observed a significant positive cross-sectional effect of antidepressant use in semantic memory. Different agents within the major class of antidepressants may affect cognitive function differently. There are several classes of antidepressants,^42^ and we did not have enough statistical power to investigate each class separately. However, in a previous study, both TCA use and SSRI use demonstrated similar risks to cognitive function in healthy individuals after controlling for self-reported depression symptoms.^5^ Moreover, the exact mechanisms of the different antidepressant classes and how they affect cognition in depressed or non-depressed individuals is not fully understood.^42^

Whereas cognitive impairments have consistently been reported in users of both benzodiazepines^21,43^ and related drugs increasing GABA transmission,^21,44,45^ the current and most other longitudinal studies did not find benzodiazepine use to be associated with cognitive decline.^24,46^ Although evidence regarding the association between benzodiazepine use and cognitive decline is conflicting, stronger links between benzodiazepine use and cognitive decline have emerged from studies investigating long-acting benzodiazepines.^46^ A longitudinal study investigating the use of sedative drugs and incident cognitive decline in older individuals did not find significant associations between use of sedatives and cognitive decline.^47^ However, we found that discontinuing anxiolytics and/or hypnotics and sedatives was significantly associated with improvement in visuospatial ability in relation to individuals continuing using these medications, and with improvement in both episodic memory and visuospatial ability in relation to individuals not using these medications. This indicates that any negative cognitive effects of using these medications may be reversed after withdrawal.

Some limitations of the present study should be acknowledged. Potential confounders that could influence the relationship between CNS medication use and cognitive task performance such as sleep problems or anxiety were not controlled for, as these data were not available. Further, a large part of individuals could not answer for how long they were using medications. Thus, data on duration of CNS mediation use were not included in the present study. Furthermore, data on self-reported medication use was limited to collection at the four time-points. However, it can be viewed as a strength that medication use was based upon participants’ reported actual medication use rather than pharmacy dispensing or records of medications prescribed to participants. Another strength of the present study is the long follow-up time of 15 years. Most importantly, our analyses to investigate the effect of CNS medication use on cognitive decline were based on CNS medication use at all four occasions rather than using baseline medication data. Individuals that discontinued the respective CNS medication during the follow-up period were excluded from these analyses, and instead included in our analyses to investigate the effect of discontinuing CNS medications on cognitive change.

In summary, although our results may be confounded by subjacent conditions, they suggest that long-term use of CNS medications may have domain-specific negative effects on cognitive performance over time, whereas the discontinuation of these medications may partly reverse these effects. These results open up for future studies that address subjacent conditions on cognition to develop a more complete understanding of the cognitive effects of CNS medications.

